# Mistake gating leads to energy and memory efficient continual learning

**DOI:** 10.64898/2026.04.16.718919

**Authors:** Aaron Pache, Mark CW van Rossum

## Abstract

Synaptic plasticity is metabolically expensive, yet animals continuously update their internal models without exhausting energy reserves. However, when artificial neural networks are trained, the network parameters are typically updated on every sample that is presented, even if the sample was classified correctly. Inspired by the human negativity bias and error-related negativity, we propose ‘memorized mistake-gated learning’—a biologically plausible plasticity rule where synaptic updates are strictly gated by current and past classification errors. This reduces the number of updates the network needs to make by 50% ∼ 80%. Mistake gating is particularly well suited in two cases: 1) For incremental learning where new knowledge is acquired on a background of pre-existing knowledge, 2) For online learning scenarios when data needs to be stored for later replay, as mistake-gating reduces storage buffer requirements. The algorithm can be implemented in a few lines of code, adds no hyper-parameters, and comes at negligible computational overhead. Learning on mistakes is an energy efficient and biologically relevant modification to commonly used learning rules that is well suited for continual learning.

---

Experimental evidence suggests that biological learning is a metabolically costly process. While the reasons are currently not fully understood, these costs can be substantial. For instance, flies subjected to aversive conditioning and subsequently starved, died 20% sooner than control flies [Mery and Kawecki, 2005]. Similarly, flies doubled sucrose consumption after long term memory formation [Plaçais and Preat, 2013], and memory can be boosted by mitochondrial stimulation [Amrapali Vishwanath et al., 2026]. This points to an evolutionary imperative to learn frugally, that is, with minimal updates to the brain’s networks [Pache and van Rossum, 2023].

Indeed, humans don’t learn equally from every sample, but place far more importance on mistakes and prediction errors, a phenomenon known as the negativity bias [Rozin and Royzman, 2001]. A familiar example is the ‘jolt’ we experience when typing a word incorrectly. When such mistakes occur, a large event-related potential known as the error-related negativity (ERN) is elicited in the EEG, which usually precedes behavioural changes like post-error slowing and post-error improvement in accuracy [Kalfaoğlu et al., 2017]. Indeed, subjects learn better on stimuli that evoke a larger EEG [de Bruijn et al., 2020]. Moreover, the ERN has been linked to dopamine signalling [Holroyd and Coles, 2002] - a plasticity modulator associated to persistent forms of synaptic plasticity (late-phase LTP) [O’Carroll and Morris, 2004].

In contrast, most training algorithms for artificial neural networks update their parameters on all samples, even if classified correctly. In particular for online continual learning, such as natural vision where the sample variation is virtually limitless, this appears excessive. First, late in learning most samples will be correctly classified and might be skipped; second, if concrete samples need to be stored for later processing, focusing on the mistakes reduces offline storage buffer requirements.

Previous work has found that for many datasets, it suffices to learn from a small subset of critical samples termed the ‘core-set’ [Agarwal et al., 2005], effectively setting the importance weight to zero for most samples. However, finding the core-set is hard and might require another model trained beforehand to identify them [Toneva et al., 2019, Wang et al., 2018, Paul et al., 2021, Coleman et al., 2020]. Alternatively, importance sampling relies more on certain samples than others during training [Katharopoulos and Fleuret, 2018]. For a new, unseen dataset, it won’t be clear what the core-set is. One ideally would have an algorithm that automatically finds important samples in an online setting, which is particularly relevant for biological and continual learning.

To better understand biological learning as well as develop more human-like AI, we explore whether mistake gating can reduce putative metabolic cost of learning and memory buffer requirements.

## Results

### Mistake-gated learning

We examine a supervised learning scenario. In such scenarios the network is trained by minimizing a surrogate loss function, commonly a mean-square error or cross-entropy loss. The reason for using a surrogate loss is that the true loss – whether the classification is correct or not – is not differentiable. While this is common practice, any sample’s surrogate loss is almost always non-zero even if classified correctly, and weights will almost always be updated. What happens if we instead make the update conditional on incorrect answers?

To test this we first train a feed-forward neural network to classify digits from the well-known MNIST dataset. We use a standard (non-convolutional) network with a single hidden layer with 200 hidden units. After learning, when presented with a test image, the output neuron with the highest activation should correspond to the class label of the presented digit. We trained the network with standard Stochastic Gradient Descent with a default learning rate of 0.01. For biological realism samples are presented one at a time, i.e. without batching (see Discussion).

A first idea is to only update the weights when a sample is misclassified. If on the other hand the sample is correctly classified, no update occurs and the algorithm skips to the next sample, as in the classic perceptron algorithm [Rosenblatt, 1958, Rimer et al., 2001, Rimer and Martinez, 2006]. We call this pure mistake-gating. With pure mistake-gating the training accuracy grows rapidly compared to traditional backprop and quickly reaches perfection, Fig. 1a (cf. red to black curves). The generalization performance, as evaluated by measuring performance on the test set, grows more slowly than traditional backprop, Fig. 1b. Yet, the number of updates steps is still substantially less than traditional backprop (black curve), Fig.1c.

**Figure 1:**
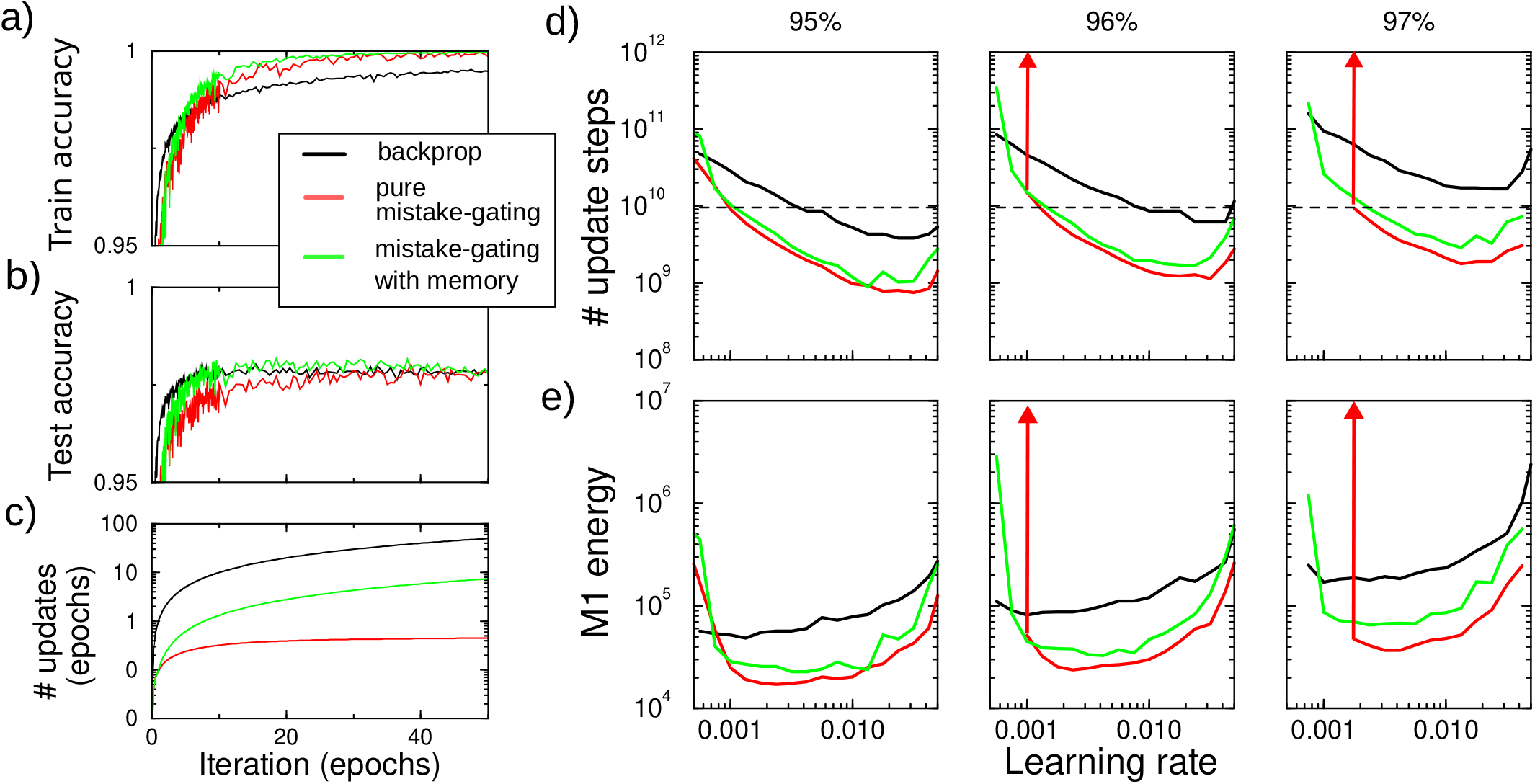
Comparison of standard back propagation rules to mistake-gated learning rules on the MNIST digit classification task. Standard backpropagation (black), pure mistake gated learning which only update when a mistake is made (red), and mistake-gated learning which updates when the sample is misclassified or was misclassified in the past (green; the focus of this study). a) Training accuracy a function of the number of training cycles. Accuracy improves more rapidly with mistake gated learning rules compared to backprop, despite using fewer updates. b) Test-set accuracy grows more slowly than training accuracy for mistake gating. c) The total number of parameter updates normalized by network size and dataset size. Backprop requires many more parameter updates than either mistake gating rule . d) The number of backpropagation steps required to reach certain test accuracy (95%, 96% and 97% percent) as a function of learning rate. An intermediate learning rate is optimal for all algorithms. Both pure mistake gated (red) and with memory (green) require far fewer training steps than backprop (black). Pure mistake-gated learning (red) can however never reach the higher accuracies, whilst with memory it can. (dashed line: number of updates over one epoch under backprop for reference). e) The putative *M*_1_ energy measure shows a similar behaviour as the number of updates, albeit at a lower optimal learning rate.

While pure mistake gated learning achieves a low error rate on the training set, the generalization on the test data is fragile. To show this we attempt to train to a certain criterion test-set accuracy. Pure mistake gating cannot reach the highest test-set accuracies when the learning rate is small (red curves in right-most panels of Fig.1d). This limits the generality and practical use of pure mistake gating. The reason is that gating results in decision boundaries close to the training samples, and hence a slight change in input can easily switch the output. Instead, traditional backprop steers the network away from the decision boundaries by updating even if the output is correct, thereby improving generalization performance on the test-set. When learning rates are large, mistake gating overshoots the decision boundary, leading to more robust solution. One way to increase robustness is to add a margin to the learning, leading to the well-known hinge loss used in support vector machines [Doğan et al., 2016]. Biologically, this requires knowledge of the activation of the neurons with respect to a particular margin and metabolically, requires knowing the margin that minimises energetic cost. Thus we concentrate on an automatic, margin-less alternative here.

### Memorized mistake gating

As pure mistake-gated learning is fragile, we modified the algorithm to memorize all incorrect samples. It updates the synapse when the sample is currently misclassified but also when it re-encounters a sample that was at *any previous time* misclassified. We call this *memorized mistake gating*. When training with this learning rule, it uses more update steps than pure mistake-gated learning, Fig. 1c (green). Yet, it is no longer fragile and still uses only a fraction of the update steps that regular backprop uses across the optimal range of learning rates and target criteria, Fig. 1d (green). Only in the limit of very small learning rates, where all samples produce effectively a random output on the first pass and so most samples are labeled as mistakes, memorized mistake gating uses as many updates as backprop.

We also measured the cumulative *L*_1_ norm of the updates, or *M*_1_ energy, that was proposed as a approximation for metabolic cost of biological learning, for instance as a anabolic/katabolic measure for the number of receptors added to and removed from the synapses [Li and van Rossum, 2020]. Mistake gating also reduces the *M*_1_ energy. This is despite that the average updates in mistake gating are on average larger than for backprop, as the incorrect samples lead to larger gradients.

Thus while memorized mistake-gating requires a similar or larger number of forward passes through the network, it drastically reduces the number of the synaptic updates required to achieve a certain performance. As this holds across learning rates (Fig.1c+d), the results can not be attributed to a change in effective learning rate. For brevity, from here on we focus on the memorized variant of mistake gating, and loosely refer to standard, ungated training as “backprop”.

Implementation of memorized mistake gating is straightforward. It requires a Boolean storage array with as many elements as there are samples in the dataset. All entries are initialized to false, but if at any time during learning a sample is misclassified, the array element for that sample is set to true and updated on immediately; it is never reset and every time that sample is encountered again, the network is updated. The mistake-gating learning rule introduces no extra parameters, and requires no tuning, the boolean array represents a negligible storage overhead, and is easily added to any existing code. We visualize each algorithm using a two-dimensional task in Fig. 2. Memorized mistake-gating yields a similar update sparsity as pure mistake-gating but since it learns on previously incorrect samples has the robust decision boundary of backprop.

**Figure 2:**
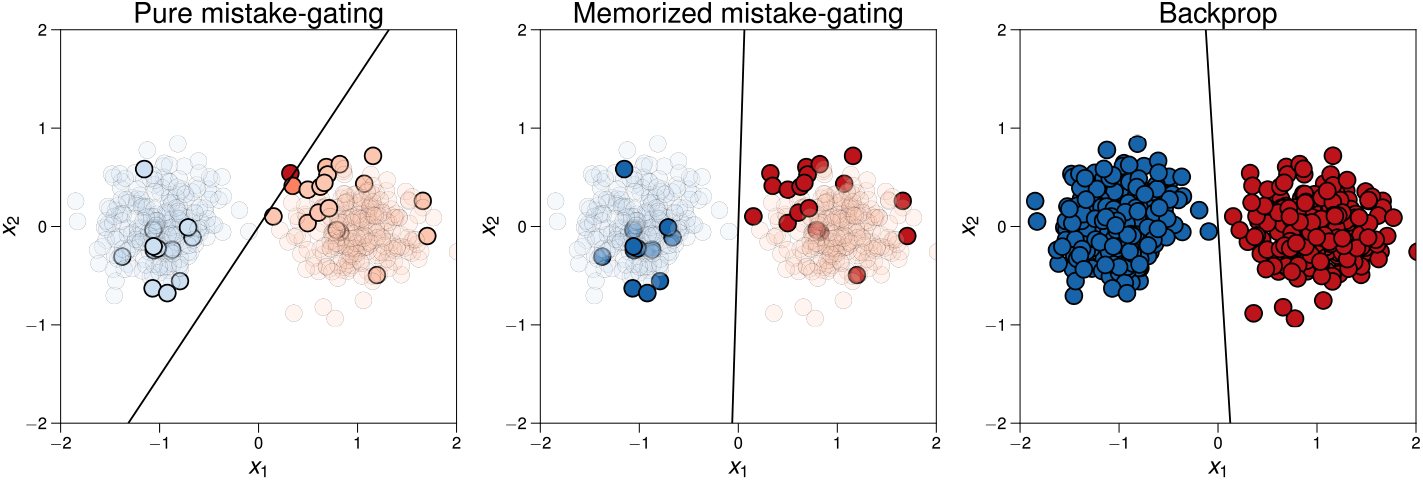
Visualization of mistake-gating on a simple 2D problem. Samples are colored according to how frequently they triggered an update and unused samples are transparent. Networks were trained for 50 epochs, with 500 samples using a small learning rate of 0.05. Pure-mistake gating concentrates updates on samples nearest the decision boundary; once a decision boundary is found, learning ceases. Memorized mistake-gating distributes updates over samples that were previously incorrect, resulting in a more robust decision boundary. Backprop uses all samples, yeilding a boundary similar to mistake-gating but without the sparse updating.

### Mistake-gating reduces memory capacity requirements for offline learning

Next, we examined how the benefits of mistake gating carry over to larger datasets. This is particularly relevant for continual learning, as most real-world data is drawn from a much larger dataset. This is most apparent in natural vision, where scene, projection and lighting changes can yield practically infinite sample variations. Including such data variation is important as it improves generalization performance, and data augmentation is therefore common practice to enlarge datasets [LeCun et al., 2015, Kaplan et al., 2020]. Yet, it also exacerbates the inefficiency of standard learning methods.

To examine the dependency on dataset size, we used the extended MNIST (EMNIST) data set, which contains 240,000 training samples, compared to 60.000 in MNIST [Cohen et al., 2017]. We trained networks on random subsets of EMNIST and tested on the full EMNIST test set (40000 samples) until it reached an accuracy of 98%.

Using standard backprop, the number of updates required to reach criterion varies only weakly with dataset size, Fig 3a. Indeed, this is what one would expect if the training subsets sufficiently cover the full data-set. Mistake gating consistently reduces it to 12%, irrespective of dataset size.

**Figure 3:**
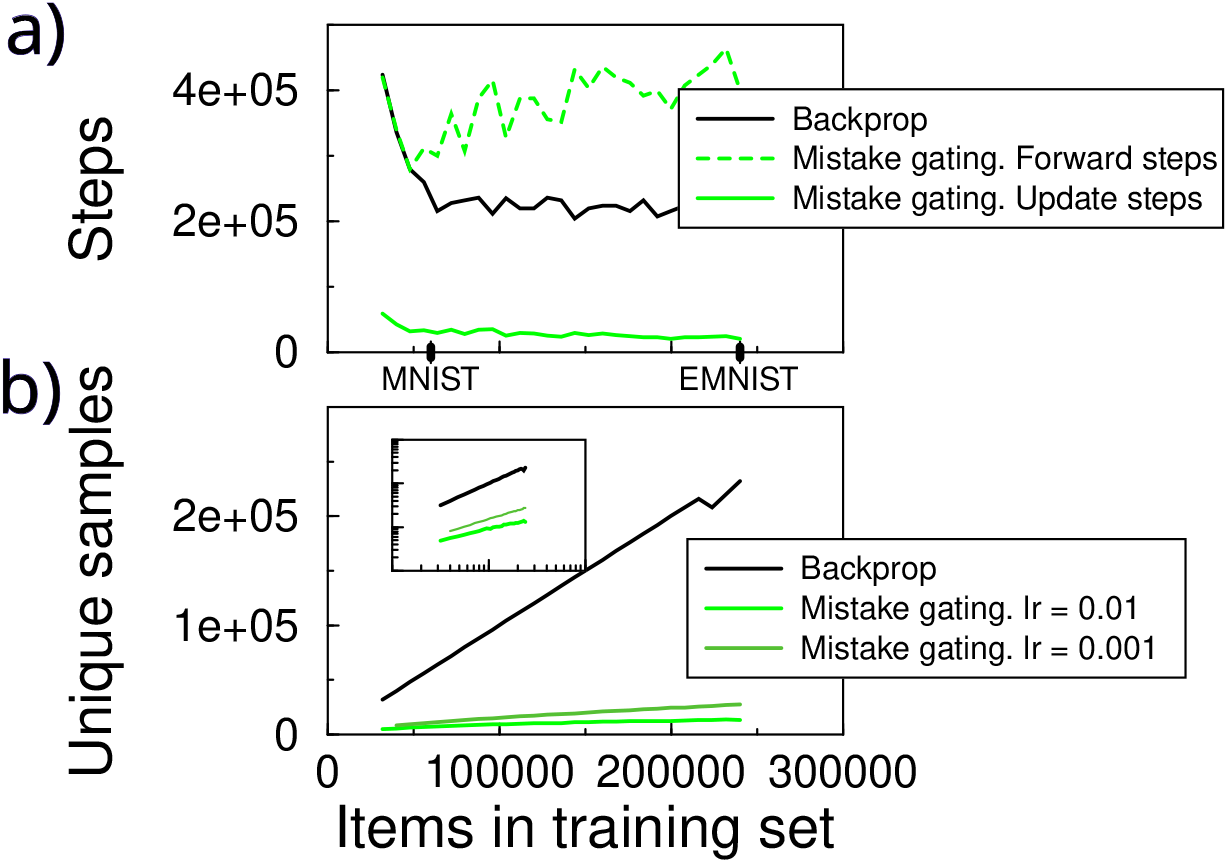
Effect of dataset size on mistake gating. The data set size was varied by using a subset of the Extended MNIST dataset. Networks were trained until they reached 98% accuracy on the EMNIST test set. Top: The number of forward steps and number of updates. Mistake gated learning requires more forward steps than backprop (dashed green), but skips over most so that fewer update steps are needed (solid green). For backprop (black) the number of update and forward steps is identical. Bottom: Mistake gated learning uses far fewer unique samples than regular backprop. The dashed line shows a powerlaw fit. Inset: same data on a log-log scale.

In addition to the number of updates, a related performance metric of mistake gating learning is the number of *unique* samples required for training, i.e. the core-set size. In machine learning it is common to have unlimited access to the dataset and algorithms cycle over all data until the required performance is reached. In biological learning, however, one typically does not have this luxury and data might need to be stored for offline replay. The hippocampus is widely considered as an example of such a temporary storage buffer, which stores events during the day, and during rest and sleep replays them, consolidating the information into cortical networks. Consistent with mistake-gated learning, evidence suggests that errors gate hippocampal memory storage [Sinclair et al., 2021]. By identifying the critical samples, mistake gating reduces the memory requirements for this temporary buffer.

To study this we assume that all information first has to be stored in a temporary buffer before it can be committed to the long-term storage. We stress that this an extremely abstracted model of consolidation, ignoring changes in representation, orthogonalization and schema extraction known to occur in the hipoocampus. We also assume the worst case scenario that no intermediate consolidation events occur, otherwise less memory would be required.

The number of unique samples needed for training grows with data-set size, Fig. 3 bottom. By construction, in regular backprop the number of unique samples is identical to the dataset size, provided that more than one training epoch is needed. Storing only incorrect samples requires however only a fraction of the storage. Interestingly, the number of unique samples that mistake-gated learning selects to update from grows sub-linearly in dataset size (light green curve). A fit yields ∼ *S*^0.51*±*0.01^, where *S* is the dataset size. When the learning rate was slowed from a default of 0.01 to 10^−3^, the exponent was 0.67*±*0.01 (dark green). In summary, while the number of update steps is approximately invariant across dataset size, the relative savings in memory requirements increase for larger datasets. Thus mistake gating allows for more efficient learning of large datasets, whether occurring naturally or augmented artificially.

### Scaling to more complex tasks

The MNIST and EMNIST tasks are easy to learn. With a properly tuned learning rate, high test accuracy on the MNIST task can already be reached using only single exposure to a part of the training dataset, i.e. within one single epoch, Fig. 1. We wondered whether the savings will diminish when tasks are more difficult and take many passes over the dataset. To test this, we first parametrically increased task difficulty by blurring the standard MNIST images with a 2D Gaussian kernel. After blurring, mean and standard deviation of the pixel intensities were matched to the original statistics. Although the blurring is in principle fully invertible, correct classification requires more precise calculations on pixel intensities and learning slows down [Ahmad, 2025]. The number of epochs required increases rapidly for both backprop and mistake-gated learning as the Gaussian kernel size increases, Fig. 4a. However, the number of update steps that memorized mistake gating requires remains at about 20% of backprop, Fig. 4a. Further analysis revealed that already in the first epoch the performance reaches ∼90%, so mistake-gating still saves. In line with this, the number of unique samples that mistake-gated learning tags as mistakes remains limited, ranging from ∼8000 (out of 60000 samples) without blurring to ∼23,000 at maximum blur (not shown).

**Figure 4:**
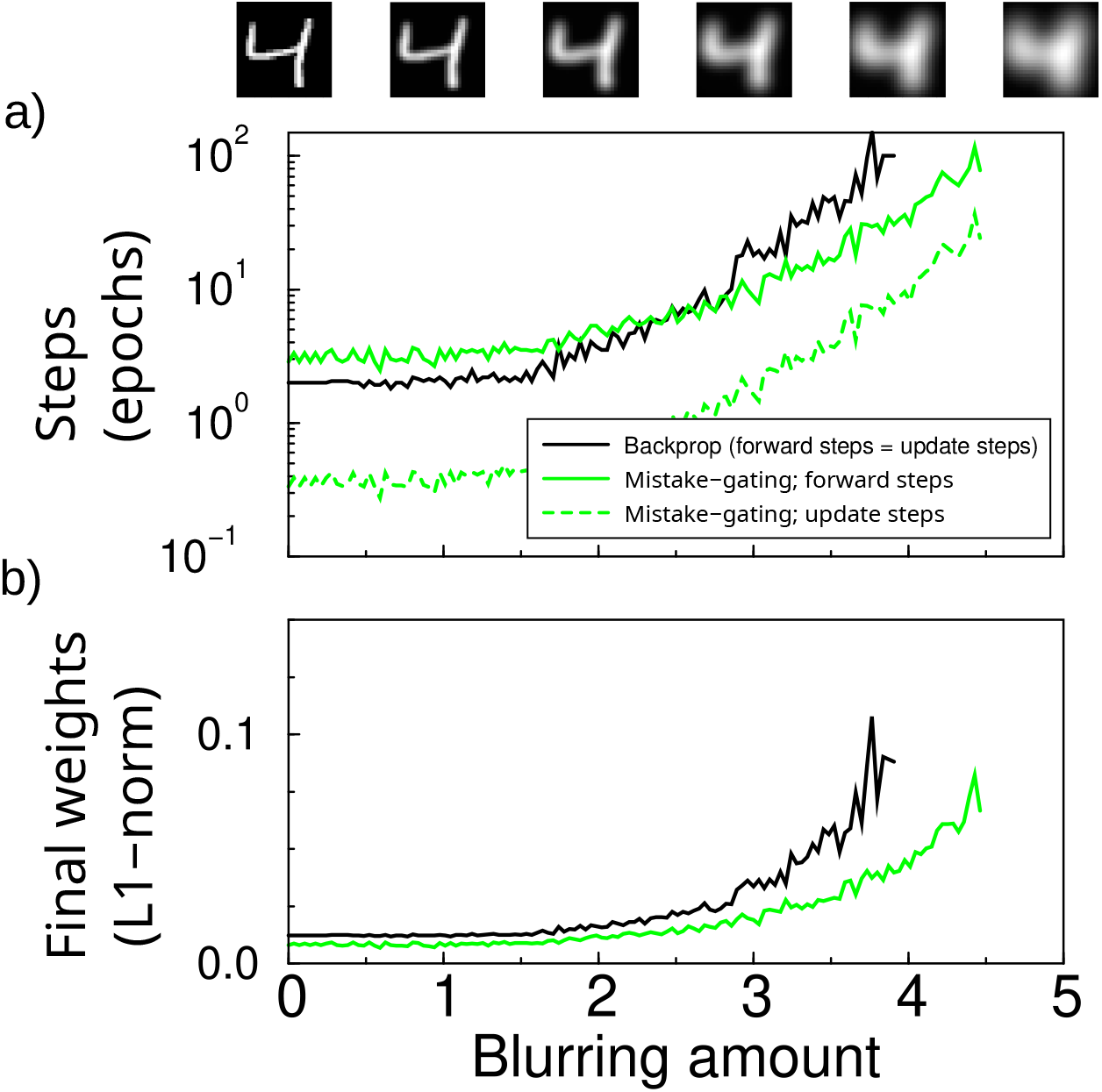
Mistake gating for dense, correlated datasets. The standard MNIST dataset was blurred with a 2D Gaussian filter, the blurring amount is parameterized as the standard deviation of the filter measured in pixels. a) The number of passes through the data set (measured in epochs) to reach 97% test accuracy increases strongly for blurred images. Note the log scale. The ratio of update steps of mistake gating learning compared to backprop is approximately constant. b) The L1 norm of the final weights (measured relative to the initial condition) pooled over the full network. Mistake-gating leads to smaller magnitude weights.

Interestingly, while mistake-gating requires more forward steps on regular MNIST, for the strongest blur the number of forward steps is actually less. The reason is hard to pin-point exactly, but backprop updates on all samples some of which become detrimental for learning under high blur; memorized mistake-gating filters these samples out becoming is less influenced by the noisy descent. Mistake gating finds final weights with smaller *L*_2_ and *L*_1_ norm, than backprop, Fig. 4b. In other words, it has a regularizing effect and we hypothesize that these smaller norms make learning in later phases easier.

Next we attempted the more challenging CIFAR-10 dataset, which contains 28×28 RGB images (60,000 train and 10,000 test) of 10 categories of animals and objects. To mimic hierarchical visual processing, we used the convolutional Densenet121 network, which has 121 convolutional layers [Huang et al., 2018].

When learning from scratch, mistake gating did not yield any savings (not shown). The reason was that almost all samples were incorrect in the first epochs, and hence are labeled as mistakes. However, biological organisms rarely learn complex tasks from a blank slate; instead, they build upon established sensory representations. We wondered if the representations that develop in the early layers could be reused for new categories. We therefore examined an incremental learning scenario. We first pre-trained the network on 7 out of 10 classes over 200 epochs (using ADAM, batching, and augmentation) to a test (training) accuracy of 96.2% (99.2%) and recorded the resulting weights. Subsequently, the remaining 3 categories were included in the dataset and training was continued on the full dataset (learning rate 0.001, single batch).

It is known that incremental learning can be faster by only allowed modification of later layers [Yosinski et al., 2014, Sorrenti et al., 2023]. Therefore we restricted the plasticity in varying amounts. Resnet-121 has 4 blocks of layers, and a final linear layer. Plasticity was either allowed in all layers; allowed in block 2 and beyond; block 3 and beyond; or, only block 4 and the output layer, Fig. 5 diagram. Batchnorm layers were always frozen.

**Figure 5:**
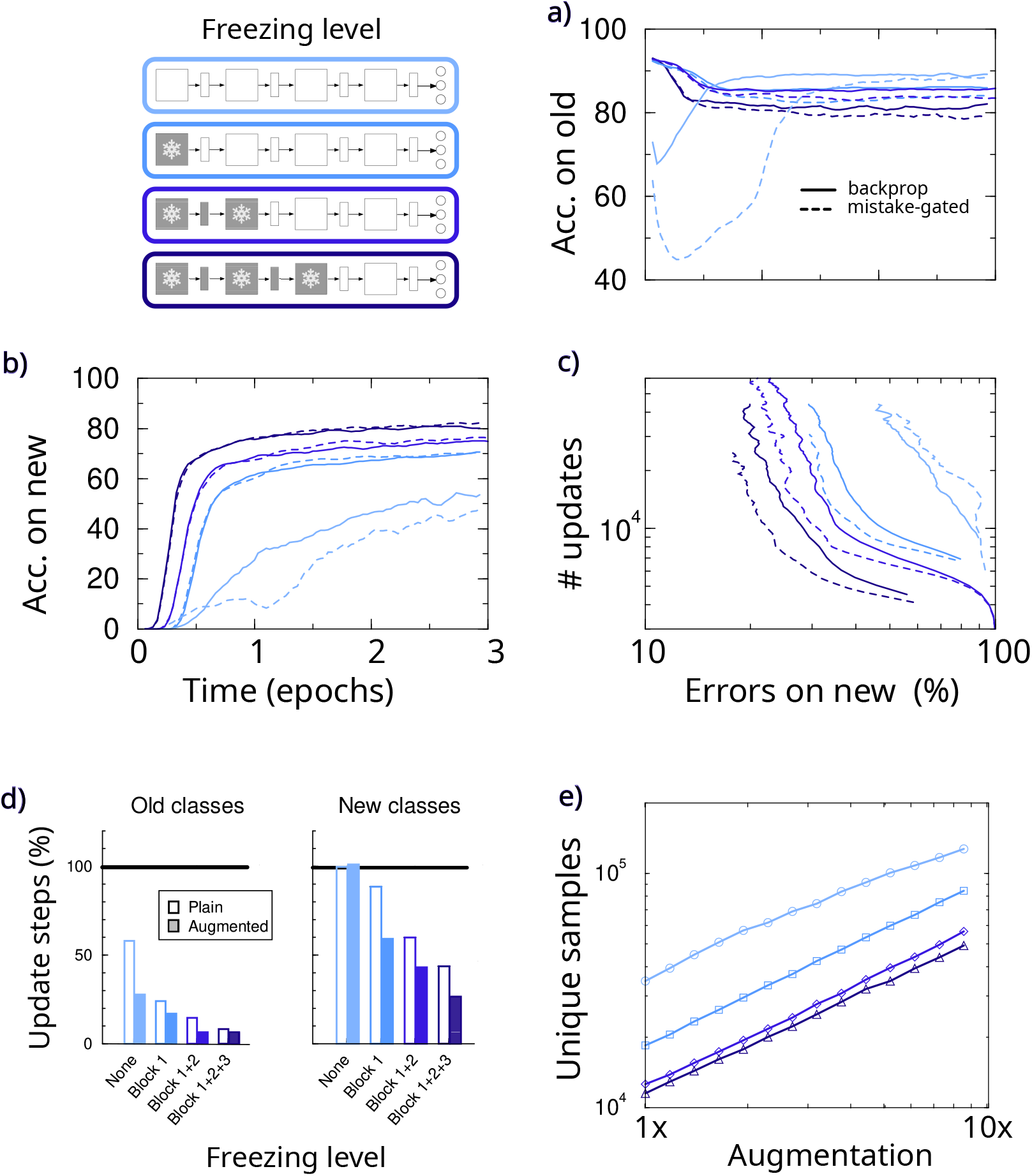
Mistake gating in incremental learning. Performance on a CIFAR-10 network, pre-trained on 7 classes after which 3 additional classes are introduced. After the pre-training the network was frozen to different degrees. Light blue: nothing frozen; darkest blue: all but last block frozen. a) Test set accuracy on old classes. The most frozen networks maintain best performance on old classes. Mistake-gating (dashed) used a similar number of forward steps (epochs) as backprop (solid curves). b) Test set accuracy on new classes, the most frozen networks learned most quickly to incorporate the new classes. c) The number of updates on the new data required to a certain test accuracy. Mistake gating (dashed curves) approximately halved the number of required updates in partly frozen networks, compared to backprop. d) Plot of saving compared to backprop vs freezing level, showing most savings for largely frozen networks, and for augmented datasets. The saving was estimated as the ratio in number of updates at the highest accuracy levels. e) The number of unique samples (old and new) required to reach 85% accuracy scaled sub-linearly with dataset size when using mistake gating.

The learning curves strongly dependent on the plasticity restrictions, Fig. 5a+b. Interestingly, when just the final block was plastic (darkest blue), learning was fast and accurate. Instead, when the network was fully plastic (lightest blue), the introduction of the new classes knocked down accuracy of the old classes and also the new classes were learned much more slowly.

What is the effect of mistake-gating in this case? Unlike above, where mistake-gating typically required more forward steps, here mistake-gating had virtually no effect on the learning curves of old or new categories; the dashed and solid curves overlapped, Fig.5a+b. However, mistake-gating uses again far less updates to reach the same accuracy level. Unsurprisingly, this is most striking for the old categories on which the network was already trained, Fig.5d. However, also for the new categories it reached similar accuracy using less updates, Fig.5c+d. We repeated this using ten-fold augmented data (mirroring and cropping). Training on augmented data led to relatively larger savings, using a smaller fraction of updates, Fig.5d (hatched bars).

Finally, we examined the number of unique samples required to train to a given test accuracy (85%) on the new classes. We changed the dataset size by sampling from the fully augmented data-set. As with the EMNIST data, we found a sub-linear power-law relation between dataset size and number of samples used, with respective exponents 0.571 (all plastic), 0.710, 0.717, 0.681 (block 4), Fig. 5e. For instance, with just the last block plastic, the number of unique memorized samples is 56 × 10^3^, vs 500 × 10^3^.

In summary, in challenging incremental learning scenarios, mistake-gating can substantially reduce the number of updates required to learn new categories provided that the networks are pre-trained. Independently, incremental learning is much quicker when plasticity is restricted to later layers (further reducing metabolic cost).

## Discussion

In summary, when training neural networks, one typically modifies the parameters for every sample. However, in biological learning plasticity is sparse, possibly because of the high associated metabolic costs to it. Mistake-gating can be seen as a temporal sparsification, automatically selecting the samples for learning in an online manner. Ultimately, the update inefficiency of standard training methods can be traced back to the fact that traditional backprop relies on a differentiable, and hence surrogate, loss function. As a result synapses are updated regardless of whether the classification is correct or incorrect. While using the surrogate loss has the benefit to steer the parameters away from the decision boundaries, and thereby improving generalization performance, it also means that every sample leads to a parameter update. This is wasteful. Eventually, late in training most networks end up in a regime where most samples are correct and mistake-gating will be highly beneficial. While earlier studies have shown that learning can indeed be limited to a subset of critical samples, most algorithms require a fixed data-set and sometimes even a double pass over the dataset, which is incompatible for the scenarios we considered.

While we concentrated on typical backpropagation learning, mistake-gating can be used with other, more biological learning algorithms [e.g. Sacramento et al., 2018]. It also naturally combines with curriculum learning [Bengio et al., 2009, Wang et al., 2022], as well as class imbalanced datasets [Johnson and Khoshgoftaar, 2019], since rare classes would be classified incorrectly more often.

In terms of experimental predictions, mistake-gating predicts that memoranda that were once incorrect but now correct, lead to additional updating. This could be tested by measuring the EEG response to such stimuli in memory experiments. We predict an error or error-like signal when such memoranda are encountered. The hippocampus has long been suggested as a temporary buffer to allow for offline learning. Mistake gated learning suggests that the content of the hippocampus is carefully titrated. On one hand, only events that evoke an error signal need to be stored. On the other hand, such events should be kept in memory, even if they do not evoke an error anymore, but did so in the past. Exactly how long the hippocampus should hold on to these patterns, is a subject for future study and will likely also depend on concurrent demands.

Related to this, it is interesting to note that traditionally pattern sparsification is known to increase hippocampal storage *capacity* [Tsodyks and Feigelman, 1988]. Our results with blurred MNIST show that sparse patterns need less storage for offline learning. Thus pattern sparsification can also reduce storage *requirements*.

One can wonder whether mistake gating benefits current machine learning. In machine learning it is common to use batching. That is, rapid GPU-based parallel computation of the gradient for many samples is followed by a single, summed update. Batching thus also reduces the number of updates. In its basic form mistake gating does not work well in batched learning, because the probability that *all* its constituent samples are correct, decreases exponentially with batch size, nullifying the benefits of lazy learning. While outside of the scope of the current paper, software engineers might be able to dynamically construct batches that only contain the mistaken sample [see also Sathiyanarayanan et al., 2025].

In biological systems, a batching-like mechanism can be implemented by including an additional, transient form of plasticity. Single sample updates are stored in transient plasticity and consolidated into late-phase plasticity only once a threshold is reached, an algorithm we termed synaptic caching [Li and van Rossum, 2020]. This saves energy because transient plasticity is metabolically much cheaper than late phase plasticity [Mery and Kawecki, 2005, Potter et al., 2009, Plaçais and Preat, 2013, Amrapali Vishwanath et al., 2026]. The slower the decay of the transient plasticity, the less consolidation events are needed. Provided that the transient memory decays only when updates are made, mistake gating reduces the number of consolidation events, compared to standard backprop with synaptic caching, thus further reducing energy needs. Future effort will be to combine the benefits of mistake learning with competitive updating which reduces the number of updates in large, over-dimensioned networks [van Rossum and Pache, 2024].

## Methods

Code is available on github.com/vanrossumlab/mistake-gating26.

## Acknowledgments

It is a pleasure to thank Claudia Danielmeier and Josh Khoo for discussion. This research was supported by a grant from NVIDIA and some simulations utilized an NVIDIA RTX A6000.

## References

P. K. Agarwal, S. Har-Peled, K. R. Varadarajan, et al. Geometric approximation via coresets. Combinatorial and computational geometry, 52(1):1–30, 2005.

N. Ahmad. Correlations are ruining your gradient descent, 2025. URL https://arxiv.org/abs/2407.10780.

A. Amrapali Vishwanath, T. Comyn, R. G. Mira, C. Brossier, C. Pascual-Caro, M. Faour, K. Boumendil, C. Chintaluri, C. Ramon-Duaso, R. Fan, et al. Mitochondrial ca2+ efflux controls neuronal metabolism and long-term memory across species. Nature Metabolism, pages 1–22, 2026.

Y. Bengio, J. Louradour, R. Collobert, and J. Weston. Curriculum learning. In Proceedings of the 26th Annual International Conference on Machine Learning, ICML 2009, pages 41–48, New York, NY,

USA, 2009. Association for Computing Machinery. ISBN 9781605585161. doi: 10.1145/1553374.1553380. URL https://doi.org/10.1145/1553374.1553380.

G. Cohen, S. Afshar, J. Tapson, and A. van Schaik. Emnist: an extension of mnist to handwritten letters, 2017. URL https://arxiv.org/abs/1702.05373.

C. Coleman, C. Yeh, S. Mussmann, B. Mirzasoleiman, P. Bailis, P. Liang, J. Leskovec, and M. Zaharia. Selection via proxy: Efficient data selection for deep learning, 2020.

E. R. de Bruijn, R. B. Mars, and R. Hester. Processing of performance errors predicts memory formation: Enhanced feedback-related negativities for corrected versus repeated errors in an associative learning paradigm. European Journal of Neuroscience, 51(3):881–890, 2020. doi: 10.1111/ejn.14566. URL https://onlinelibrary.wiley.com/doi/abs/10.1111/ejn.14566.

Ü. Doğan, T. Glasmachers, and C. Igel. A unified view on multi-class support vector classification. Journal of Machine Learning Research, 17(45):1–32, 2016.

C. B. Holroyd and M. G. H. Coles. The neural basis of human error processing: reinforcement learning, dopamine, and the error-related negativity. Psychol Rev, 109(4):679–709, Oct. 2002.

G. Huang, Z. Liu, L. van der Maaten, and K. Q. Weinberger. Densely connected convolutional networks, 2018. URL https://arxiv.org/abs/1608.06993.

J. M. Johnson and T. M. Khoshgoftaar. Survey on deep learning with class imbalance. Journal of Big Data, 6(1):27, Mar 2019. ISSN 2196-1115. doi: 10.1186/s40537-019-0192-5. URL https://doi.org/10.1186/s40537-019-0192-5.

Ç. Kalfaoğlu, T. Stafford, and E. Milne. Frontal theta band oscillations predict error correction and posterror slowing in typing. J Exp Psychol Hum Percept Perform, 44(1):69–88, Apr. 2017.

J. Kaplan, S. McCandlish, T. Henighan, T. B. Brown, B. Chess, R. Child, S. Gray, A. Radford, J. Wu, and D. Amodei. Scaling laws for neural language models. arXiv preprint arXiv:2001.08361, 2020.

A. Katharopoulos and F. Fleuret. Not all samples are created equal: Deep learning with importance sampling. In J. Dy and A. Krause, editors, Proceedings of the 35th International Conference on Machine Learning, volume 80 of Proceedings of Machine Learning Research, pages 2525–2534. PMLR, 10–15 Jul 2018. URL https://proceedings.mlr.press/v80/katharopoulos18a.html.

Y. LeCun, Y. Bengio, and G. Hinton. Deep learning. Nature, 521(7553):436–444, 2015. ISSN 0028-0836. doi: 10.1038/nature14539. Insight.

H. Li and M. C. W. van Rossum. Energy efficient synaptic plasticity. Elife, 9:e50804, 2020.

F. Mery and T. J. Kawecki. A cost of long-term memory in drosophila. Science, 308(5725):1148, May 2005. doi: 10.1126/science.1111331.

C. M. O’Carroll and R. G. Morris. Heterosynaptic co-activation of glutamatergic and dopaminergic afferents is required to induce persistent long-term potentiation. Neuropharmacology, 47(3):324–332, sep 2004. doi: 10.1016/j.neuropharm.2004.04.005.

A. Pache and M. C. van Rossum. Energetically efficient learning in neuronal networks. Current Opinion in Neurobiology, 83:102779, 2023.

M. Paul, S. Ganguli, and G. K. Dziugaite. Deep learning on a data diet: Finding important examples early in training, 2021.

P.-Y. Plaçais and T. Preat. To favor survival under food shortage, the brain disables costly memory. Science, 339(6118):440–442, 2013. ISSN 0036-8075. doi: 10.1126/science.1226018.

P.-Y. Plaçais and T. Preat. To favor survival under food shortage, the brain disables costly memory. Science, 339(6118):440–442, 2013.

W. Potter, K. O’riordan, D. Barnett, S. Osting, C. Burger, and A. Roopra. Metabolic regulation of hippocampal ltp via the energy sensor ampk: Psm06-03. J. of Neurochemistry, 108:100–101, 2009.

M. Rimer and T. Martinez. Classification-based objective functions. Machine Learning, 63(2):183–205, May 2006. ISSN 1573-0565. doi: 10.1007/s10994-006-6266-6.

M. Rimer, T. Andersen, and T. Martinez. Speed training: improving the rate of backpropagation learning through stochastic sample presentation. In IJCNN’01. International Joint Conference on Neural Networks. Proceedings (Cat. No.01CH37222), volume 4, pages 2661–2666 vol.4, 2001. doi: 10.1109/IJCNN.2001.938790.

F. Rosenblatt. The perceptron: A probabilistic model for information storage and organization in the brainnformation storage and organization in the brain. Psychological Review, 65(6):386–408, 1958.

P. Rozin and E. B. Royzman. Negativity bias, negativity dominance, and contagion. Personality and Social Psychology Review, 5(4):296–320, 2001. doi: 10.1207/S15327957PSPR0504\_2.

J. Sacramento, R. P. Costa, Y. Bengio, and W. Senn. Dendritic cortical microcircuits approximate the backpropagation algorithm. In Advances in neural information processing systems, pages 8721–8732, 2018.

S. M. Sathiyanarayanan, X. Hao, S. Hou, Y. Lu, L. Sevilla-Lara, A. Arnab, and S. N. Gowda. Progressive data dropout: An embarrassingly simple approach to faster training, 2025. URL https://arxiv.org/abs/2505.22342.

A. H. Sinclair, G. M. Manalili, I. K. Brunec, R. A. Adcock, and M. D. Barense. Prediction errors disrupt hippocampal representations and update episodic memories. Proceedings of the National Academy of Sciences, 118(51):e2117625118, 2021.

A. Sorrenti, G. Bellitto, F. P. Salanitri, M. Pennisi, C. Spampinato, and S. Palazzo. Selective freezing for efficient continual learning. In Proceedings of the IEEE/CVF International Conference on Computer Vision, pages 3550–3559, 2023.

M. Toneva, A. Sordoni, R. T. des Combes, A. Trischler, Y. Bengio, and G. J. Gordon. An empirical study of example forgetting during deep neural network learning, 2019.

M. V. Tsodyks and M. V. Feigelman. The enhanced storage capacity in neural networks with low activity level. Europhys Lett, 6:101–105, 1988.

M. C. van Rossum and A. Pache. Competitive plasticity to reduce the energetic costs of learning. PLOS Computational Biology, 20(10):e1012553, 2024.

T. Wang, J. Huan, and B. Li. Data dropout: Optimizing training data for convolutional neural networks. In 2018 IEEE 30th International Conference on Tools with Artificial Intelligence (ICTAI), pages 39–46, 2018. doi: 10.1109/ICTAI.2018.00017.

X. Wang, Y. Chen, and W. Zhu. A survey on curriculum learning. IEEE Transactions on Pattern Analysis and Machine Intelligence, 44(9):4555–4576, 2022. doi: 10.1109/TPAMI.2021.3069908.

J. Yosinski, J. Clune, Y. Bengio, and H. Lipson. How transferable are features in deep neural networks? Advances in neural information processing systems, 27, 2014.

